# FibrilPaints for visualisation and ubiquitination of Huntingtin amyloid fibrils

**DOI:** 10.64898/2025.12.19.695423

**Authors:** Françoise A. Dekker, Emile van Weert, Guy Mayer, Tommaso Garfagnini, Júlia Aragonès Pedrola, Alfred C.O. Vertegaal, Assaf Friedler, Stefan G. D. Rüdiger

## Abstract

Huntington’s disease (HD) is caused by expansion of a polyglutamine tract in the huntingtin (Htt) protein, leading to aggregation of the exon 1 fragment (HttEx1) into amyloid fibrils. HttEx1 forms one of the lowest-complexity amyloid cores known, its fibril core consists of a single amino acid, glutamine. With emerging therapies improving patients’ prospects by silencing expression of HTT, tools to monitor HttEx1 aggregation become essential for timely intervention and next-generation therapeutics. Here, we show that the peptide FibrilPaint1 selectively binds HttEx1Q44 fibrils without interacting with monomeric protein, allowing to measure and trace HttEx1 amyloid fibrils. Using the FibrilRuler assay, we tracked fibril formation from early species to larger clustered assemblies. The non-fluorescent variant, FibrilPaint20, was used to recruit the E3 ubiquitin ligase CHIP to HttEx1 fibrils, enabling site-specific ubiquitin tagging. However, unlike Tau fibrils, ubiquitinated HttEx1 fibrils resisted proteasomal degradation. This reveals a fundamental difference in how amyloids with extremely low–complexity cores respond to cellular clearance machinery. Together, our findings establish the FibrilPaint peptide family as a toolset for the detection and molecular targeting of amyloids, providing new opportunities to study protein aggregation and act as building blocks for future diagnostic and therapeutic strategies in neurodegenerative diseases.

**Highlights:**

- FibrilPaint1 selectively binds HttEx1Q44 amyloid fibrils and allows monitoring of fibril growth using the hydrodynamic radius (FibrilRuler).
- FibrilPaint20 recruits the E3 ligase CHIP to Htt fibrils, enabling ubiquitination.
- Despite successful ubiquitination, Htt fibrils resist proteasomal degradation in vitro, highlighting structural barriers.
- FibrilPaint provides a scaffold for functional targeting of amyloids with diagnostic and therapeutic potential.

**Graphical abstract:** Graphical abstract
The FibrilRuler Test: FibrilPaint enables measurement of Huntingtin fibril size during aggregation
After a short lag-phase following removal of the protective MBP tag by Factor Xa, fibrillation proceeds rapidly. Subsequent fibril clustering further accelerates growth, leading to exponential increases in aggregate size.

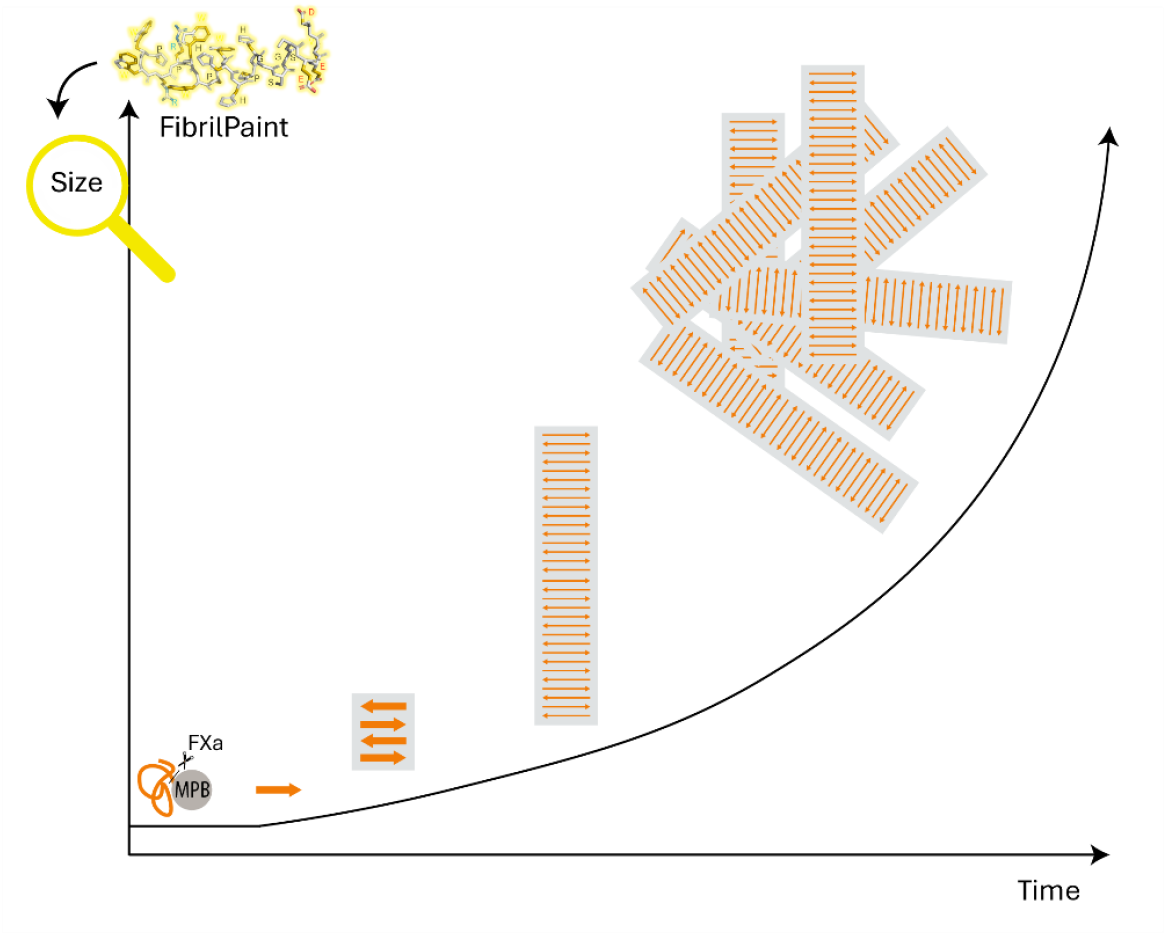

## Introduction

Huntington’s disease (HD) is an autosomal dominant neurodegenerative disorder characterized by progressive tissue loss in the neocortex and striatum, leading to motor, cognitive, and psychiatric symptoms [1, 2]. It affects approximately 6 - 9 individuals per 100,000 in the Western world [3]. The disease is caused by an expanded CAG trinucleotide repeat in the Huntingtin (HTT) gene, which encodes an abnormally long polyglutamine (polyQ) tract in exon 1 of the huntingtin protein (Htt) [1, 4-7]. A polyQ stretch exceeding 35 glutamines is pathogenic, and disease onset and severity correlate with its length [2, 8].

mHttEx1 forms amyloid fibrils that accumulate as large intracellular inclusions and resist cellular clearance mechanisms [9, 10]. Structurally, the HttEx1 fragment consists of three domains: an N-terminal region, the polyQ tract, and a C-terminal segment. The polyQ core forms the amyloid backbone, assembling into antiparallel, non-twisting β-sheets [11, 12]. The N-terminal region initiates intermolecular contact, and together with the C-terminus, forms a dynamic “fuzzy coat” that surrounds the fibril core [12-14]. This stands in contrast to other disease-associated amyloids, such as Tau, α-synuclein, and Aβ, which possess chemically diverse, multi-residue cores capable of twisting and forming multiple conformations [12, 15].

Therapeutic strategies are emerging to target or prevent the aggregation process [16-18]. Notably, uniQure recently reported encouraging results from two Phase I/II clinical trials (NCT04120493, NCT05430117) using AMT-130, a gene therapy designed to silence HTT expression [19]. High-dose treatment resulted in up to 75% slowing of clinical decline in early-manifest HD patients, alongside reductions in cerebrospinal fluid neurofilament light (NfL), a marker of neurodegeneration [19]. These findings represent a breakthrough and provide strong support for therapeutic approaches that prevent or delay amyloid formation. However, gene silencing alone does not eliminate pre-existing aggregates, underscoring the need for complementary strategies and tools to track residual pathology.

To this end, we need reliable methods for detecting and quantifying mHttEx1 species [20-22]. Genetic testing and symptom presentation are effective for diagnosis of HD, and monitoring can be supported by indirect biomarkers such as NfL levels or MRI-based measures of brain atrophy [23, 24]. While novel PET tracers like [^11C]CHDI-180R show promise for imaging mHTT aggregates, they are still limited by high intersubject variability and low signal-to-background ratios [25, 26]. So, huntingtin pathology remains undetectable in patients. Chemically, this poses an interesting challenge, because the polyQ core lacks the chemical diversity typically required for selective molecular recognition [12, 15].

We previously developed FibrilPaints: modular peptides that selectively bind amyloid fibrils and can be functionalized for detection or enzymatic recruitment of the ubiquitin system [27-29]. Although all variants affected Tau aggregation kinetics, only FibrilPaint1 bound with sufficient affinity to enable fibril size quantification by Flow-Induced Dispersion Analysis (FIDA), a setup we termed the FibrilRuler test [27]. In this study, we examined their effect on Huntingtin Exon 1 with 44 glutamines (HttEx1Q44), the fragment that aggregates in HD. We also explore targeted modification of fibrils via FP20-mediated recruitment of the ubiquitination machinery. Our approach presents a modular toolkit for studying Htt aggregation.

## Results

### FibrilPaint1 specifically binds HttEx1Q44 fibrils, not monomers

We first assessed the effect of FibrilPaint addition on HttEx1 aggregation, both to establish a baseline for binding detection and to enable direct comparison with our earlier findings in Tau [27]. Aggregation was initiated by cleavage of the N-terminal maltose-binding protein (MBP) tag using Factor Xa (FXa), which triggered fibril formation **(Fig. 1A)**. Aggregation kinetics were monitored by Thioflavin T (ThT) fluorescence, an established reporter of cross-β amyloid structures [30]. FibrilPaint1–4 were added at final concentrations of 0.2, 1, and 2 µM, and ThT fluorescence was recorded over 10 h **(Table 1)**. All peptides decreased the final ThT plateau of 20 µM HttEx1Q44 aggregation in a concentration-dependent manner, indicating interaction with fibril formation or competitive binding with ThT. This effect was less pronounced than previously observed for TauRD [27]. Among the four peptides, FibrilPaint4 showed the lowering of the ThT platform to 15 ± 8 %, followed by FibrilPaint2 (57 ± 8 %), FibrilPaint3 (68 ± 13 %), and FibrilPaint1 (79 ± 17%).

**Table 1.**
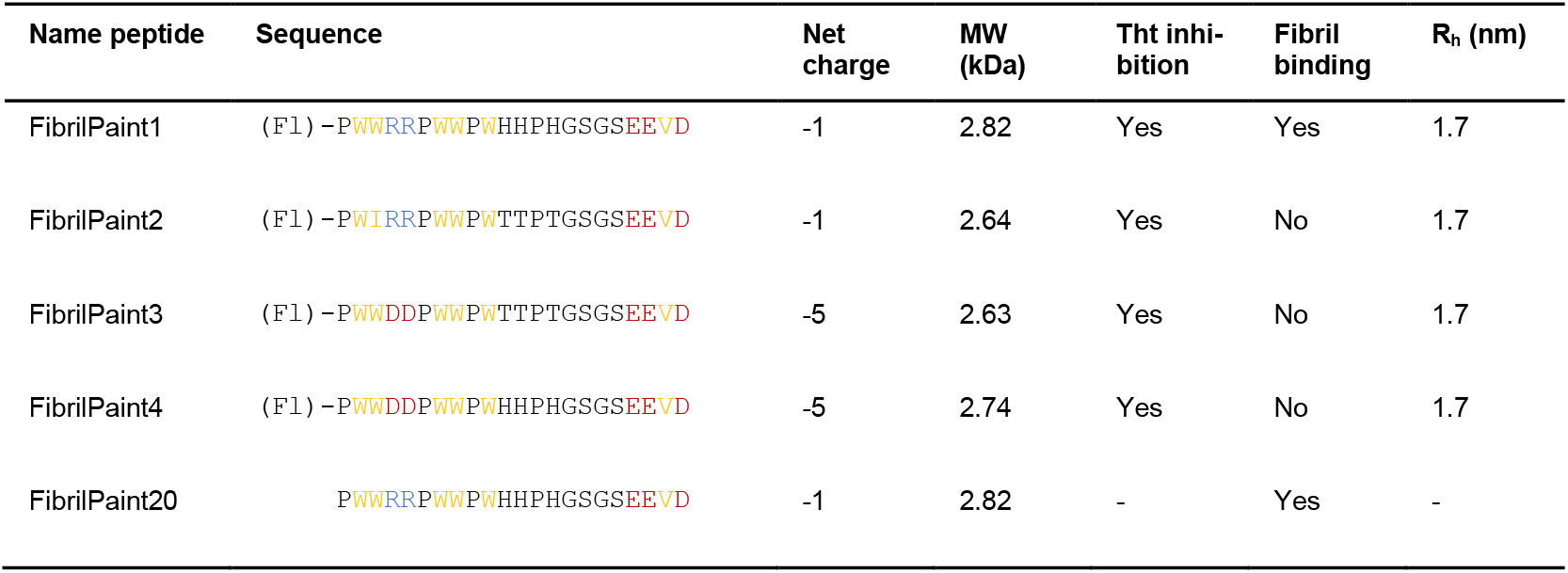
Peptide names and sequences. (Fl) indicates an N-terminal fluorescein label, which was excluded from calculations of molecular weight and amino acid length. Amino acids are coloured according to the YRB scheme (hydrophobic residues in yellow, negatively charged residues in red, and positively charged residues in blue) (1). Net charge refers to the cumulative charge of each peptide at physiological pH. Fibril binding was assessed by detecting Tau amyloid fibrils using Flow-Induced Dispersion Analysis (FIDA). The hydrodynamic radius (R_h_) corresponds to the particle size measured by FIDA in the absence of binding interactions.

**Figure 1.**
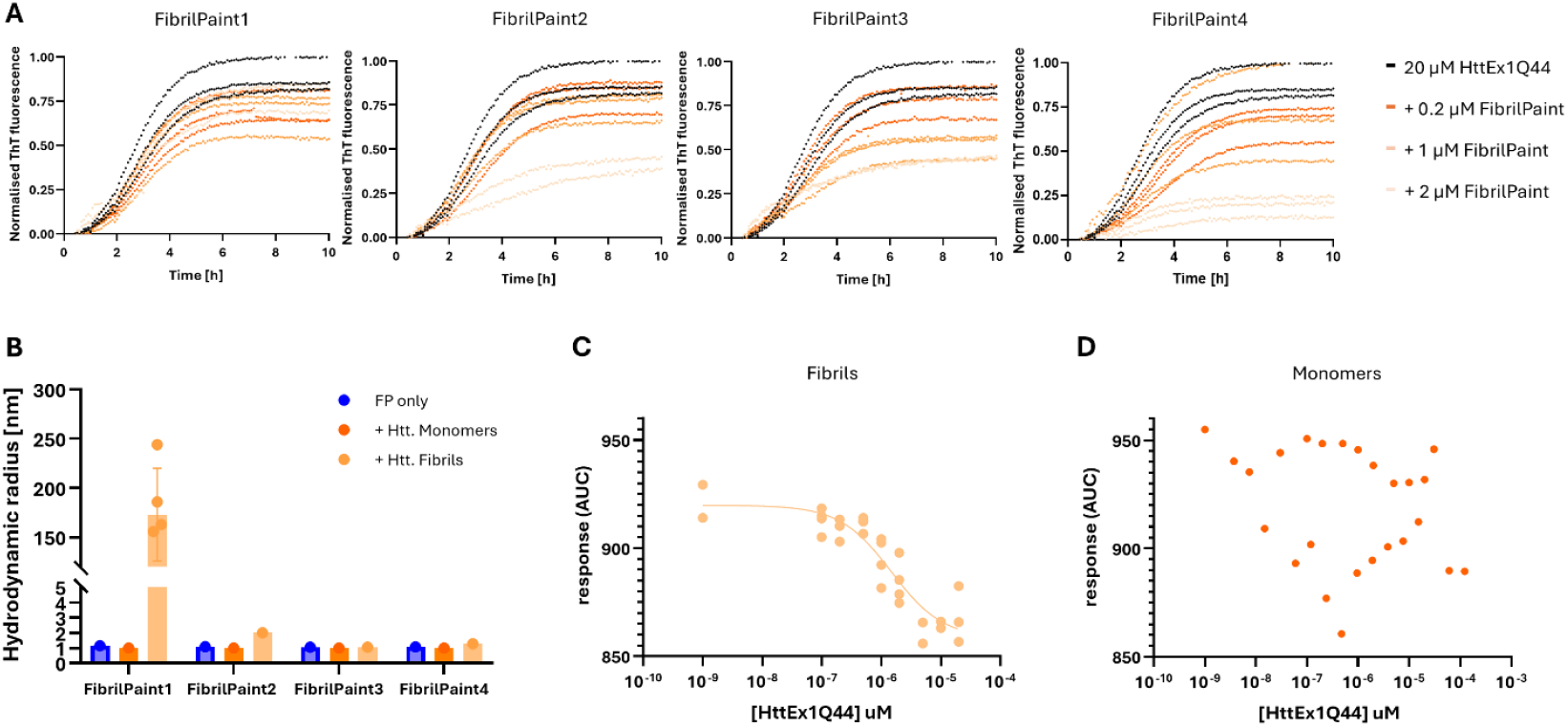
FibrilPaint1 specifically binds Huntingtin Exon 1 (HttEx1Q44) amyloid fibrils. **(A)** Thioflavin T (ThT) fluorescence assay showing the effect of FibrilPaint1–4 on the aggregation kinetics of 20 µM HttEx1Q44 following MBP-tag cleavage by Factor Xa (FXa). Representative graph with all triplicates shown from N=3. Fluorescence intensity is normalised to the maximum value measured at the aggregation plateau, indicating peptide-dependent reduction of the plateau value. **(B)** Hydrodynamic radius (R_h_) determined by Flow-Induced Dispersion Analysis (FIDA) for FibrilPaint1–4 alone (blue), in the presence of HttEx1Q44 monomers (dark orange), or after incubation with 20 µM preformed HttEx1Q44 fibrils (light orange, 5 h aggregation). Binding to fibrils results in a pronounced R_h_ increase. All individual measurements shown. **(C, D)** Microscale thermophoresis (MST) analysis of 50nM FibrilPaint1 interaction with up to 150 µM HttEx1Q44 left to aggregate for 5-hours **(C)** or measured immediately (monomers) **(D)**. MST traces were recorded using a 1.5 s on-time at medium infrared power. Apparent dissociation constants (Kd ± SD) from triplicate measurements was 1.5 µM (logK_d_ = −5.83 ± 0.28) for fibrils and not applicable (>10^-4^) for monomers.

Next, we examined the binding of FibrilPaint peptides to HttEx1Q44 amyloid fibrils using the FibrilRuler set-up [27]. HttEx1Q44 was aggregating for 5 h, corresponding to the plateau-phase of aggregation **(Fig. 1A)**, and subsequently incubated with 200 nM FibrilPaint at a final fibril concentration of 2 µM. The hydrodynamic radius (R_h_) of FibrilPaint was monitored by Flow-Induced Dispersion Analysis (FIDA) **(Fig. 1B)** [31, 32]. The average R_h_ of FibrilPaint1 increased from 1.7 ± 0.2 nm (free peptide) to 173 ± 44 nm in the presence of HttEx1Q44 fibrils, whereas no change was observed with monomeric HttEx1Q44. This demonstrates that FibrilPaint1 binds specifically to the fibrillar form of HttEx1Q44 but not to the monomeric protein. In contrast, FibrilPaint2–4 showed no detectable R_h_ increase in FIDA and were therefore ineffective in binding HttEx1Q44 fibrils **(Fig. 1B)**. These results indicate that aggregation pathways can be modulated through weak or transient interactions, rather than high-affinity binding[33-35]. Binding specificity of FibrilPaint1 was further validated by microscale thermophoresis (MST) using fluorescently labelled FibrilPaint1 (50 nM) titrated with increasing concentrations of HttEx1Q44 fibrils or monomers (0–200 µM) **(Fig. 1C, D)**. MST analysis revealed a dose-dependent interaction between FibrilPaint1 and amyloid fibrils, but no measurable binding to monomers **(Fig. 1C, D)**. The apparent dissociation constant (Kd) for fibril binding was 1.5 µM (logK_d_ = -5.83 ± 0.28), calculated assuming monomer-equivalent concentrations; because each fibril comprises multiple monomeric units, the true affinity of FibrilPaint1 for the fibrillar species is likely higher. These results establish FibrilPaint1 as a unique peptide probe that selectively binds and fluorescently labels amyloid fibrils while discriminating against the monomeric form.

### Fibril kinetics reveal continued fibril growth after ThT plateau

Next, we applied the FibrilRuler method to monitor the kinetics of HttEx1Q44 aggregation and compared its performance with conventional aggregation assays. To this end, we measured the time-dependent aggregation process in parallel using ThT fluorescence, dynamic light scattering (DLS) for size analysis, TEM imaging and the FibrilRuler test **(Fig. 2)**. DLS determines the cumulant hydrodynamic radius of particles in homogeneous solution. However, for heterogenous solutions like fibrils, it enables detection of size changes in the entire sample, but not specific changes in fibril length, which is why we refer to the readout as the apparent hydrodynamic radius (R_h_, app) [36-40]. For this reason, we applied DLS primarily to monitor trends in aggregation dynamics, rather than to determine absolute fibril sizes, as previously described for Tau aggregation [29].

**Figure 2.**
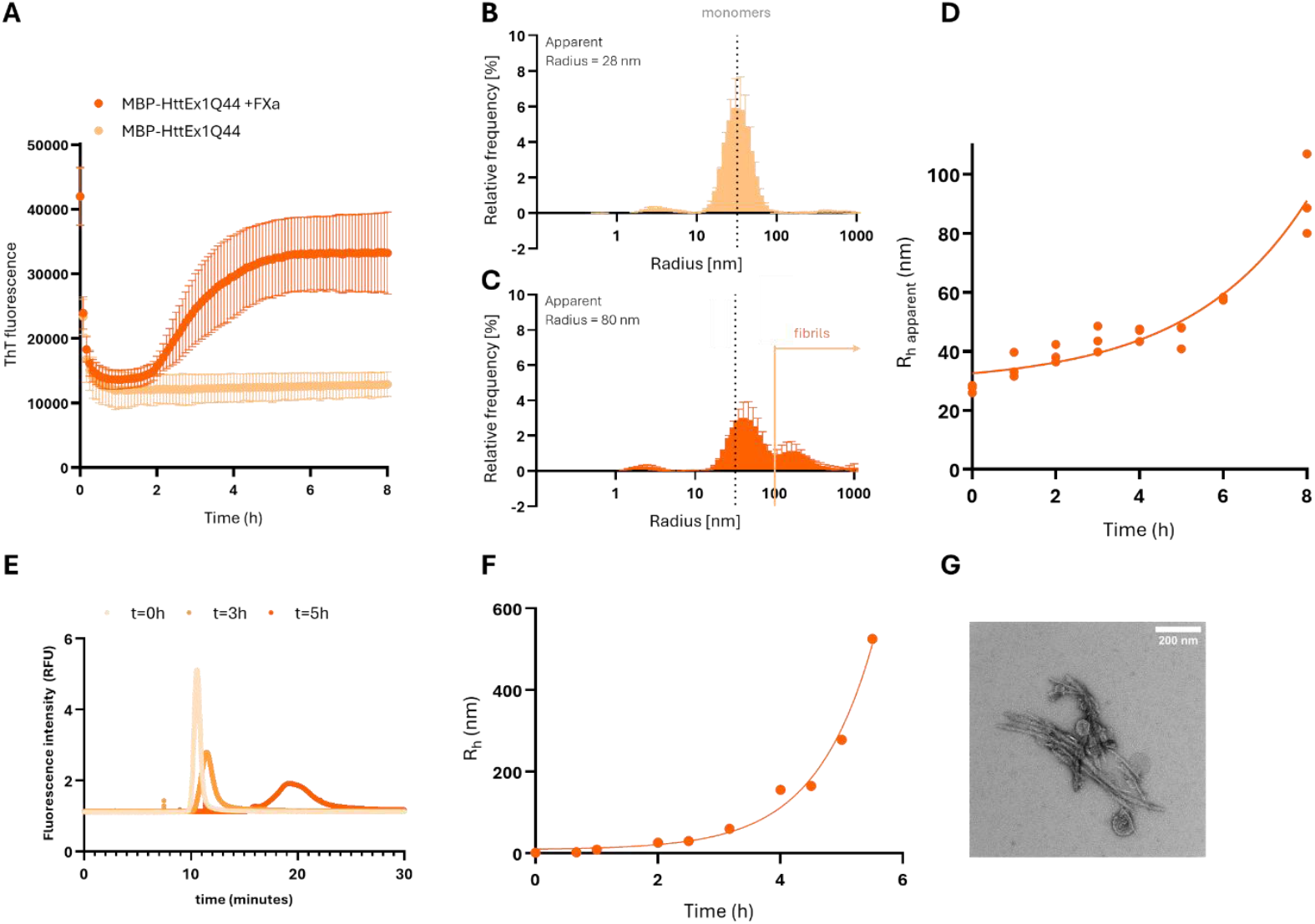
Aggregation kinetics and characterization of HttEx1Q44 fibril formation. **(A)** Aggregation of 20 µM HttEx1Q44 monitored by Thioflavin T (ThT) fluorescence in the presence (dark orange) or absence (light orange) of Factor Xa (FXa) for cleavage of the N-terminal MBP tag. Aggregation proceeds only after tag removal. **(B–C)** Representative DLS measurements of the apparent hydrodynamic radius (R_h_, app) of **(B)** monomeric MBP– HttEx1Q44 (28 nm) and **(C)** HttEx1Q44 fibrils after 8 h of aggregation (80 nm). **(D)** Time-resolved evolution of R_h_, app during HttEx1Q44 aggregation as measured by DLS. **(E)** Representative graphs of HttEx1Q44 samples at 0, 3, and 5 h using the FibrilRuler setup: using fluorescently labelled FibrilPaint1to label amyloids and subsequent FIDA analysis. Progressive delay and broadening of the FIDA peaks indicate increasing particle size, reflected in a larger hydrodynamic radius (R_h_) **(F)** Time-resolved evolution of R_h_, during HttEx1Q44 aggregation using the FibrilRuler setup. **(G)** Transmission electron microscopy (TEM) images of HttEx1Q44 fibrils after 6 h aggregation.

As a benchmark, we used the ThT assay for monitoring amyloid kinetics. The ThT fluorescence profile revealed a lag phase of 1.4 ± 0.68 h, followed by a growth phase that reached a plateau after 4.4 ± 0.35 h of aggregation, with a half-time of 2.8 ± 0.13 h **(Fig. 2A)**. The control sample containing uncleaved MBP–HttEx1Q44 (without FXa) showed no increase in ThT fluorescence, confirming that aggregation was triggered by MBP cleavage. Variations in aggregation onset between replicates reflect the spontaneous and heterogeneous nature of amyloid nucleation.

In DLS, monomeric MBP–HttEx1Q44 displayed an R_h__app of 28 nm, which is exactly 8-times larger than the 3.5 nm expected based on its molecular weight alone **(Fig. 2B)**. Previously, some MBP fusions with other proteins have been shown to form dimers, tetramers, or larger oligomers due to the oligomerization of the fused protein [41-44]. For our research purposes, it suffices to know that these species are not amyloid aggregates **(Fig. 2A)**.

Following addition of Factor Xa, the protective MBP tag is cleaved, enabling aggregation to initiate. Consistent with this, DLS measurements revealed the emergence of new peaks at larger hydrodynamic radii, accompanied by a decrease in the relative intensity of the monomer peak **(Fig. 2B, C)**. These changes indicate the formation of larger species. Over time, aggregation progressed further and with it the R_h__app further increases, reaching approximately 80 nm after 8 h **(Fig. 2C)**.

Time-resolved DLS measurements revealed an overall trend broadly consistent with ThT fluorescence during the early phase of aggregation **(Fig. 2A, D)**. The R_h__app increased gradually from 0 to 3 h, followed by a plateau-like region spanning approximately 3–5 h. From 5 h onward, R_h__app increased more strongly, exceeding 80 nm at the 8 h time point. This behaviour indicates continued growth of HttEx1Q44 aggregates beyond the initial β-sheet– forming phase and provides a complementary, label-free readout of aggregation. Notably, a monomer-associated peak remained detectable throughout the experiment, indicating that MBP-cleavage was not complete by 8 h.

Next, we analysed the aggregation kinetics of HttEx1Q44 using the FibrilRuler test. In Flow-Induced Dispersion Analysis (FIDA), the diffusion of fluorescent particles is monitored as they pass through a detector; larger particles diffuse more slowly, resulting in broader peaks and an increased R_h_. As an example, the peaks for t=0, t=3 and t=5h of 20 µM HttEx1Q44 aggregation are given **(Fig. 2E)**. Immediately after initiating aggregation (t=0 h), the R_h_ measured for HttEx1Q44 was 1.7 nm, corresponding to the signal of free FibrilPaint. T=3h already showed a clear broadening and rightward shift of the FIDA peak, consistent with progressive formation of larger assemblies, and this trend is exceeded by timepoint t=5h **(Fig. 2E)**.

Time-resolved FibrilRuler measurements revealed a continuous increase in R_h_ during aggregation **(Fig. 2F)**. After 1 h, R_h_ reached 9.4 nm, exceeding the predicted R_h_ of 3.4 nm for the unfolded monomeric HttEx1Q44 structure, indicating the presence of multimeric species. Aggregation continued to increase exponentially, reaching an average R_h_ of approximately 500 nm after 5.5 h **(Fig. 2F)**. As the particle size approached the upper limit of FIDA measurement, the final plateau value could not be determined.

The discrepancy between DLS and FIDA measurements can be attributed to the substantial fraction of monomeric protein that remains in solution. In DLS, the cumulant-derived hydrodynamic radius represents an intensity-weighted average over all species present, such that residual monomers bias the apparent size toward smaller values. In contrast, FIDA selectively reports on fibrillar species, as FibrilPaint1 binds specifically to aggregates, rendering the readout insensitive to the monomeric population.

Transmission electron microscopy (TEM) at 6h revealed that HttEx1Q44 fibrils began to form clusters **(Fig. 2G)**. HttEx1Q44 fibrils formed heterogeneous clusters of varying sizes, ranging from small aggregates composed of a few fibrils (50–100 nm) to large assemblies several micrometres in width. A typical clustered structure measured approximately 800 nm in length and 200 nm in width. Assuming a spherical geometry, this corresponds to an R_h_ of 400 nm, consistent with the 300–500 nm R_h_ values obtained from FIDA measurements.

Taken together, the results from ThT fluorescence, DLS, FIDA, and TEM consistently describe the aggregation of HttEx1Q44. The continued increase in size observed in both DLS and FIDA reflects progressive fibril growth and clustering, explaining the rapid and continuous rise detected by both methods. Although the general trend was apparent in DLS, measurement with FIDA captured absolute values of these clusters better. Such clustering likely does not generate additional ThT-binding sites, which explains why ThT fluorescence plateaus while FIDA and DLS continue to report increasing particle size. Together, these complementary assays demonstrate that HttEx1Q44 aggregation proceeds through the formation and coalescence of fibril clusters, leading to large assemblies that may remain ThT-silent but are readily detected by hydrodynamic methods **(Fig. 2A-G)**.

### FibrilPaint1 inhibits fibril clustering

Next, we analysed the effect of FibrilPaint1 when present during HttEx1Q44 aggregation, as its presence appeared to lower the ThT fluorescence plateau **(Fig. 1A)**. This reduction could result either from (1) an interaction of FibrilPaint1 with intermediate or fibrillar species, thereby altering the aggregation pathway, or (2) competition with ThT for binding sites on the fibril surface. To assess the underlying mechanism, we monitored the aggregation process using the FibrilRuler test to determine the size of the resulting assemblies.

Samples from 20 µM HttEx1Q44 aggregation reactions containing 2 µM FibrilPaint1 were collected at multiple time points and analysed by FIDA **(Fig. 3A)**. The R_h_ remained stable at approximately 28 nm throughout the 8 h reaction. Negative-stain TEM corroborated this observation, revealing fibrils that were more spatially dispersed and showed reduced bundling compared to untreated samples **(Fig. 3B)**.

**Figure 3.**
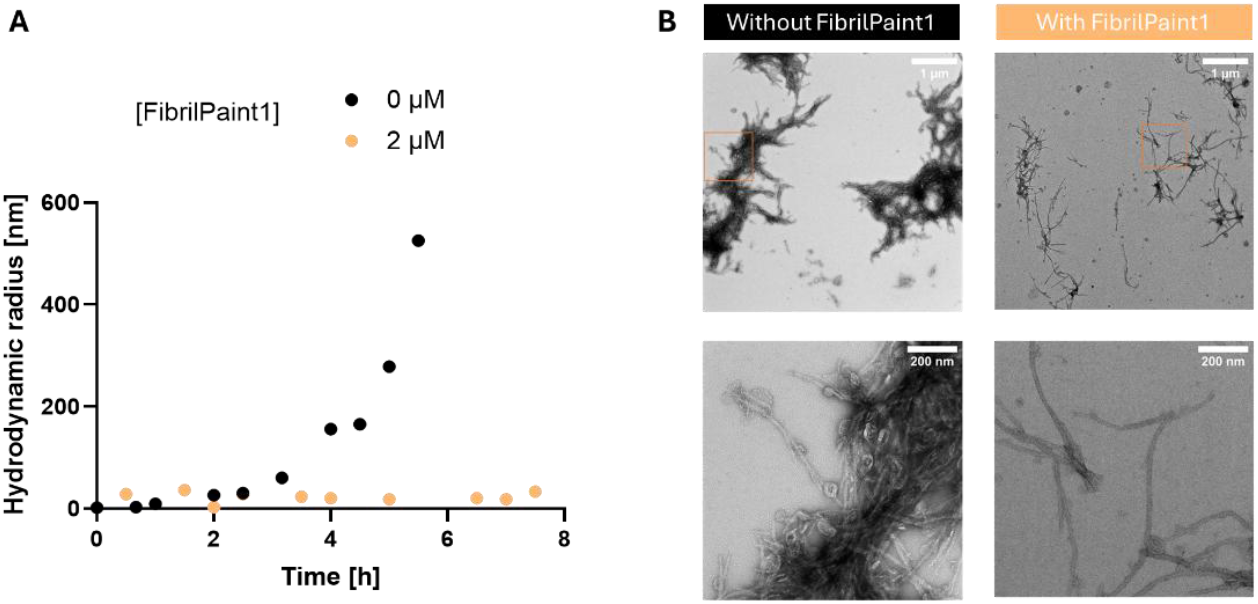
FibrilPaint interacts with HttEx1Q44 fibrils during aggregation. **(A)** Fibrils incubated with FibrilPaint1 were analysed using the FibrilRuler setup. The hydrodynamic radius (R_h_) remained approximately 28 nm throughout the experiment (light orange). For comparison, previously acquired data of HttEx1Q44 aggregation without FibrilPaint are shown alongside (black).**(B)** Representative TEM images of HttEx1Q44 fibrils grown for 6 h without (left) or in the presence of FibrilPaint1 (right). The upper panel shows a low-magnification overview (scale bar, 1 µm), while the lower panel presents a magnified view highlighting fibril morphology (scale bar, 200 nm).

Importantly, for elongated fibrillar particles, changes in axial length have only a limited influence on the measured hydrodynamic radius, as rotational tumbling in solution leads to an effective R_h_ that is dominated by the fibril cross-section rather than its length [31, 45]. In contrast, lateral association and bundling substantially increase the effective cross-sectional area and therefore result in pronounced increases in R_h_ and size dispersity. The observation that R_h_ remains constant while fibrils are clearly present indicates that fibril elongation can still occur, but that secondary processes such as lateral association or clustering are suppressed.

The reduced size dispersity observed by FIDA, together with the more isolated fibril morphology seen by TEM, is therefore consistent with FibrilPaint1 binding along the lateral surfaces of HttEx1Q44 fibrils. Such surface binding would sterically hinder close side-by-side packing of fibrils, thereby limiting bundling and preventing the formation of larger clustered assemblies, while not necessarily interfering with axial fibril elongation. This mechanism explains both the constant R_h_ and the decreased ThT fluorescence observed in the presence of FibrilPaint1.

### Conversion of R_h_ to fibril length

To better interpret the relationship between the R_h_ and fibril length, we analysed the properties of HttEx1Q44 layers based on previously published coordinates [12]. In this model, each layer of the fibril consists of four HttEx1Q44 monomers arranged into β-hairpins that stack to form a cross-β structure **(Fig. 4A, B)**. The flexible N- and C-terminal regions extend outward, forming a fuzzy coat. By stacking successive layers, we established a correlation between fibril length and the number of layers, and consequently with R_h_ values obtained by FIDA **(Fig. 4C)**. For instance, the R_h_ of 9.4 nm measured after 1 h of aggregation corresponds to a structure comprising approximately 75 layers, which are roughly 300 monomers **(Fig. 2F, 4C)**. Naturally, once lateral association occurs, this simplified single-fibril model no longer applies.

**Figure 4.**
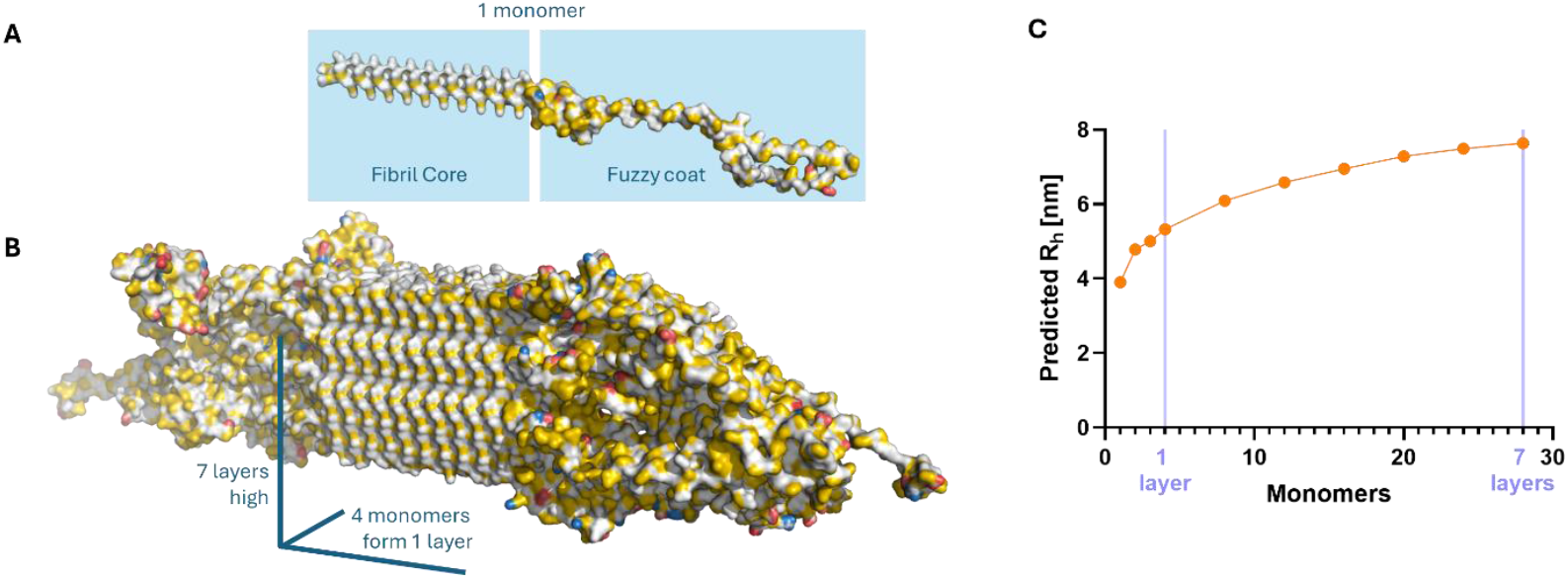
Structural and mathematical representation of Huntingtin Exon 1 (HttEx1Q44) amyloid fibrils. **(A)** Schematic representation of a single HttEx1Q44 monomer folded into its amyloid conformation, highlighting the ordered fibril core and the surrounding disordered fuzzy coat. **(B)** Model of a seven-layer HttEx1Q44 amyloid fibril reconstructed according to the NMR coordinates provided by the van der Well laboratory [12]. Models are coloured according to the YRB (Yellow-Red-Blue) scheme to indicate hydrophobic (yellow), negatively charged (red), and positively charged (blue) residues [46] **(C)** Mathematical correlation between the number of incorporated HttEx1Q44 monomers and predicted hydrodynamic radii (R_h_) in nm. Fibril layers were generated in PyMOL and their corresponding R_h_ were calculated using FIDAbio software.

After the initial layers are incorporated, fibril length approaches a linear correlation to R_h_. If the 28 nm R_h_ measured for HttEx1Q44 in the presence of FibrilPaint1 represents unbundled fibrils, the average corresponding fibril length can be estimated 570 fibril layers, over 2000 monomers. Each layer has typical interstrand spacing for β-sheets of 0.47–0.5 nm, resulting in a length of approximately 300 nm. Thus, if the fibrils are singular, we can easily convert the R_h_ with the structural coordinates to the fibril length.

### FibrilPaint can be used for ubiquitination, but subsequent proteasomal degradation does not occur

Having established that FibrilPaint1 interacts with HttEx1Q44 fibrils without promoting further aggregation, we next investigated whether a related peptide, FibrilPaint20 (FP20), could be used to actively modify these fibrils through ubiquitination [29]. FP20 shares the same sequence as FibrilPaint1 but lacks the fluorescent N-terminal tag. Both have an EEVD motif at the end, for targeting E3-ligase CHIP.

We first examined whether FP20 enhances the recruitment of the E3 ubiquitin ligase CHIP to HttEx1Q44 fibrils. FP20 was titrated at increasing concentrations against Atto-565–labelled CHIP (Fl-CHIP) **(Fig. 5A)**. The fluorescence baseline remained constant, indicating that binding did not alter fluorophore conformation [29]. Titration of HttEx1Q44 fibrils shows a dose-dependent increase in signal, which was enlarged in the presence of FP20, although the apparent dissociation constant (K_app) remained constant: 4.6 × 10^−7^ M (logK = -6.3 ± 0.7) without FP20 and to 1.7 × 10^−7^ M M (logK = -6.8 ± 0.5) in its presence **(Fig. 5A)**. This indicates that FP20 does not alter the intrinsic affinity of CHIP for fibrils, but instead increases the extent of CHIP recruitment per fibril, consistent with formation of a higher-order ternary complex.

**Figure 5.**
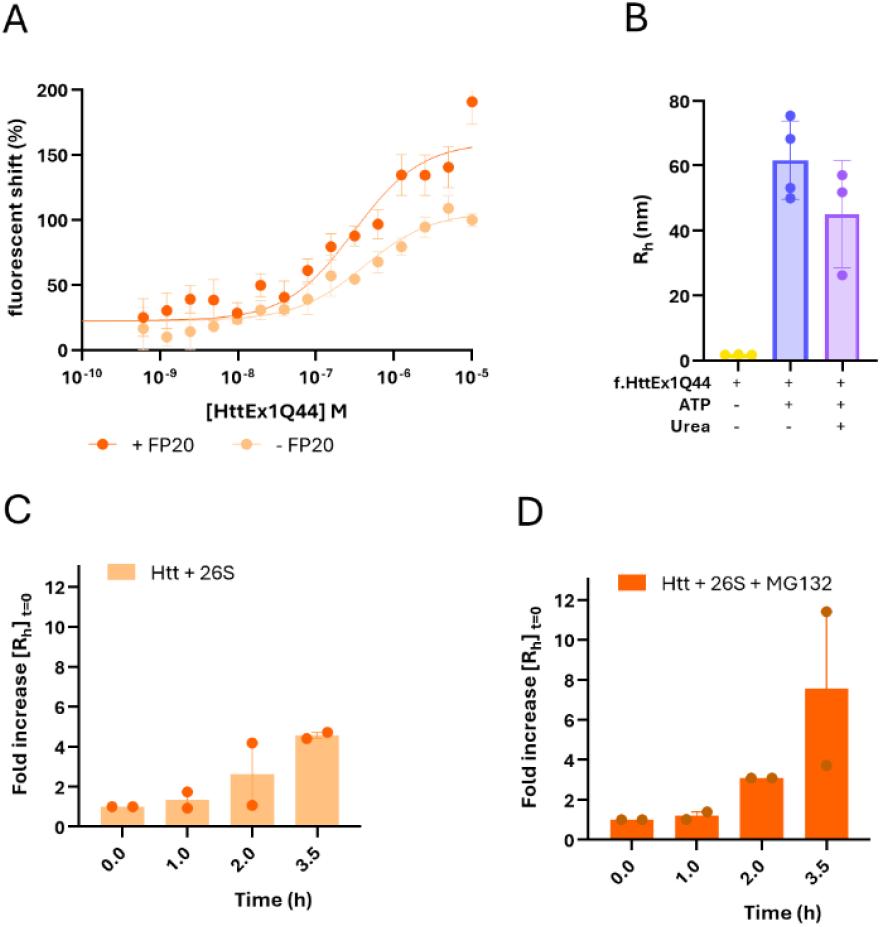
Targeting HttEx1Q44 amyloid fibrils for ubiquitination with FibrilPaint20. **(A)** Fluorescent shift of Atto565-labelled CHIP in presence of up to 100 µM HttEx1Q44 fibrils (light orange) measured with microscale thermophoresis. In the presence of FibrilPaint20 (FP20, dark orange), fluorescence shift increases for CHIP binding to HttEx1Q44 fibrils. Affinities are 4.6 × 10^−7^ M (logK = -6.3 ± 0.7) without FP20 and to 1.7 × 10^−7^ M (logK = -6.8 ± 0.5). **(B)** Ubiquitin transfer assay using fluorescently labelled ubiquitin (Fl-Ub) over the complete ubiquitination cascade (E1–E2–E3) and FP20 to HttEx1Q4 fibrils in presence of absence of ATP. In presence of ATP, the hydrodynamic radius (R_h_) increases to approximately 62 ± 12 nm. After urea treatment, the R_h_ is 45 ± 16 nm, confirming covalent attachment of ubiquitin to HttEx1Q44 fibrils. **(C)** Ubiquitinated HttEx1Q44 fibrils remain show a continued increase in size to 3.6 ± 0.2 fold the Rh,app of timepoint 0, indicating that ubiquitin tagging alone does not result in proteasomal degradation. **(D)** In the presence of proteasomal inhibitor MG132, grow in 3.5 h to 6.5 ± 5.4 increase in Rh, app.

Next, we evaluated ubiquitin transfer to amyloid fibrils using the complete in vitro ubiquitination cascade with FP20 as a recruiter. In the absence of ATP, the R_h_ was 1.8 ± 0.1 nm, corresponding to free Fl-Ub. Upon activation of the system, the R_h_ increased to 62 ± 12 nm for HttEx1Q44 fibrils, confirming ubiquitin conjugation **(Fig. 5B)**. After urea treatment, the R_h_ decreased to 45 ± 16 nm, likely due partial dissociation of fibril bundles or structural heterogeneity within the HttEx1Q44 aggregates [13, 47, 48].

Next, we assessed proteasomal degradation of HttEx1Q44 fibrils using the established DLS setup (Fig. 4D, G; Fig. 5). We purified human 26S proteasomes from HEK293 GP2 cells stably expressing hRpn11-HTBH and divided them into two aliquots for immediate use. One aliquot was preincubated for 1.5 h with 50 µM of the proteasome inhibitor MG132, while the other was supplemented with buffer. Subsequently, approximately 0.2 nM of 26S proteasome was added to each sample with ubiquitinated HttEx1Q44 fibrils [49].

We incubated HttEx1Q44 fibrils with 26S proteasomes for 3.5 h in the presence or absence of MG132. To correct for variability between fibril preparations, all data were normalized to the apparent hydrodynamic radius (R_h_, app) at the time of 26S addition, ensuring that subsequent changes reflected proteasomal activity rather than sample heterogeneity.

After 3.5 h, fibrils incubated in the presence of MG132 continued to increase in size, showing a 6.5 ± 5.4-fold increase in R_h_,app, compared to a 3.6 ± 0.2-fold increase in the absence of the inhibitor. Despite this apparent difference, the large variability, particularly in the MG132 condition, meant that the effect was not statistically significant.

Consistent with this observation, only modest and variable changes were detected at earlier time points. In the absence of MG132, R_h_,app increased to 1.33 ± 0.57 after 1 h and 2.6 ± 2.2 after 2 h, while in the presence of MG132 the corresponding values were 1.20 ± 0.36 at 1 h and 3.0 ± 0.1 at 2 h. Across all time points, the differences between conditions fell within experimental variability and did not reach statistical significance.

Taken together, these data indicate that proteasomal activity does not lead to a statistically relevant difference of fibril growth under these conditions. While a trend toward increased fibril expansion in the presence of MG132 is observed, the effects are small and highly variable, suggesting that proteasome-mediated modulation of fibril size is limited in this assay. This is in contrast to our previous findings for Tau fibrils [29].

Altogether, these findings show that FibrilPaint20 enables selective ubiquitination of HttEx1Q44 fibrils, while active proteasomes might restrict, but do not degrade, these aggregates.

## Discussion

In this study, we establish FibrilPaint1 as a tool to investigate amyloid formation by Huntingtin Exon 1 (HttEx1). We demonstrate that FibrilPaint1 binds selectively to HttEx1 fibrils, while showing no binding to monomeric species. Using the FibrilRuler assay, we reveal fibril growth ranging from early species up to large, higher-order aggregates exceeding 600 nm in radius, resulting from processes such as fibril bundling or clustering. In parallel, we show that the non-fluorescent variant FibrilPaint20 can recruit the E3 ligase CHIP to HttEx1 amyloid fibrils, enabling ubiquitination. Together, these findings establish the FibrilPaint peptide family as both a detection platform and a modular scaffold for enzymatic modification of HttEx1 fibrils.

With the binding of FibrilPaint peptides now extended beyond tauopathies to Htt, despite the lack of sequence or structural homology, it is likely that these peptides recognise a shared structural motif across amyloid fibrils [27, 50]. During HttEx1 aggregation, monomeric HttEx1 likely transitions from an α-helical or disordered conformation into a tetrameric intermediate by interactions between the N17 domains [14, 51, 52]. Upon formation of this nucleus, a β-sheet conformation can be established, and subsequent fibril elongation proceeds primarily as a monomer-dependent process [51, 52]. FibrilPaint1 does not inhibit HttEx1 aggregation, but decreased fibril clustering. This supports a model in which binding occurs at the sides, allowing new monomers to stack the fibrils.

Interestingly, while Tau fibrils tagged with FibrilPaint20 were efficiently ubiquitinated and degraded by the proteasome in prior work, HttEx1Q44 fibrils are not [29]. Explanations could be in the structural differences, which may affect the ability of the proteasome to cleave ubiquitinated fibrils. Clustering of HttEx1 fibrils may sterically hinder proteasomal engagement [53, 54]. The low complexity core presents uninterrupted stretches of glutamine–glutamine hydrogen bonds that pack into highly regular β-sheets, which may too densely stacked to disrupt [12, 54]. Intermediate Htt aggregates may actively impair proteasomal function by engaging the proteasome in a non-productive manner, potentially preventing gate opening or substrate translocation, and larger structures may trap the 26S proteasomes [53, 55].

Alternatively, ubiquitination levels may not be sufficient to enable proteasomal degradation of Htt fibrils. The placement of the ubiquitin group is likely on the fuzzy coat, as lysine residues are located within the N17 flanking region, which is also where patient-derived Htt fibrils are ubiquitinated [56]. Following ubiquitin recognition, the 19S regulatory subunits of the proteasome require access to an unstructured region to initiate substrate unfolding and translocation into the 20S catalytic core [57, 58]. For small fibrils, the proteasome can use these regions to pull the fibril apart [59]. For larger fibrils, the proteasome only degrades the fuzzy coat or the fibril is fragmented [60, 61]. This suggests that early and smaller fibrils may present more viable targets for proteasome-mediated clearance. Hence, it would be key for therapeutic success to start treatment at the earliest phase of the disease.

For HD, compounds like FibrilPaint could serve as tools to identify and characterize the specific amyloid species present under different physiological and pathological conditions. Misfolded Htt species capable of seeding aggregation have been detected in brain, cerebrospinal fluid, and even blood, yet their structural identity remains unknown [21, 24, 62-65]. Better molecular characterization of these species could help understand their toxicity and origin, and offers a gateway to targeted intervention. More broadly, approaches that recognize conserved amyloid structures rather than disease-specific sequences may enable a unified framework for studying protein aggregation across neurodegenerative disorders. As disease-modifying therapies emerge, such tools will be increasingly important for linking molecular pathology to diagnosis, monitoring, and therapeutic strategy.

## Acknowledgements

SGDR was supported by grants of the Campaign Team Huntington and Alzheimer Nederland (No. WE.03-2019-03) and a ZonMW TOP grant (No. 91215084). SGDR and ACOV are principal investigators of the Gravitation Consortium “FLOW” (024.006.036), funded by the Dutch Ministry of Education, Culture, and Science (OCW). FAD and SGDR were supported by NWO TakeOff1 (22073) and BioTech Booster (BIOB25014), funded by the Dutch Ministry of Education, Culture, and Science (OCW). AF thanks The Minerva Center for Bio-Hybrid complex systems and the Saerree K. and Louis P. Fiedler Chair in Chemistry. Measurements on the FIDA1, CLARIOstar® Plus and Monolith were done at the Protein Research Centre of Utrecht University.

## Author contributions

- Conceptualization FD JAP
- Data curation FD
- Formal analysis FD TG
- Funding acquisition SGDR
- Investigation FD EW TG GM
- Methodology FD JAP TG GM
- Supervision SGDR AF
- Visualization FD
- Writing – original draft FD
- Writing – review and editing FD SGDR TG ACOV AF

## Conflict of interest

FAD, TG, JAP, AF and SGDR are named as inventors in a patent (EP23194706, ‘Peptides for the detection of amyloid fibril Aggregates’) filed by Universiteit Utrecht Holding BV describing the peptides mentioned in this manuscript. The other authors declare no competing interests.

## Materials and methods

### Peptide synthesis and purification

Peptides were synthesized on a Liberty Blue microwave-assisted peptide synthesizer (CEM) using standard Fmoc solid-phase chemistry with Oxyma/DIC as coupling reagents. Peptide concentrations were determined by UV absorbance. N-terminal labeling was performed using 5(6)-carboxyfluorescein. Cleavage from the resin was carried out for 3 h at room temperature using a mixture of 95% (v/v) trifluoroacetic acid (TFA), 2.5% (v/v) triisopropylsilane (TIS), and 2.5% (v/v) triple-distilled water with continuous agitation. Following cleavage, solvent volume was reduced under a nitrogen stream and peptides were precipitated by addition of four volumes of cold diethyl ether (−20 °C). Samples were incubated at −20 °C for 30 min, centrifuged, and washed three times with diethyl ether before drying under nitrogen.

Dried peptides were dissolved in a 1:2 (v/v) mixture of acetonitrile and water, snap-frozen in liquid nitrogen, and lyophilized. Purification was performed using reverse-phase preparative HPLC (WATERS) on a C18 column with an acetonitrile/water gradient. Peptide identity and purity were confirmed by electrospray ionization mass spectrometry and analytical reverse-phase HPLC on a C8 column.

### HttEx1Q44 expression, purification, and fibril preparation

HttEx1Q44 was expressed in E. coli BL21 Rosetta 2 (Novagen) as a fusion protein containing an N-terminal maltose-binding protein (MBP) tag and a C-terminal His_6_ tag. Bacterial cultures were induced, harvested, and lysed following standard protocols. Clarified lysates were filtered through 0.22 µm polypropylene filters and purified using an ÄKTA chromatography system. Proteins were captured on a POROS 20MC affinity column equilibrated in 50 mM HEPES-KOH (pH 8.5) and 100 mM KCl, and eluted using a linear imidazole gradient (0–100%, 0.5 M imidazole over 10 column volumes, 30 mM NaCl).

Fractions containing HttEx1Q44 were pooled, buffer-exchanged into 50 mM HEPES and 150 mM NaCl, and concentrated using centrifugal filters (MWCO 30 kDa). Protein concentration was measured by UV spectroscopy (NanoDrop™ OneC), and purity was assessed by SDS– PAGE. Aliquots were stored at −80 °C.

Fibril formation was initiated by cleavage of the MBP tag using Factor Xa (0.1 µM) added to 20 µM HttEx1Q44 in 25 mM HEPES-KOH (pH 7.4), 75 mM KCl, 75 mM NaCl, and protease inhibitor (½ tablet per 50 mL). Samples were incubated at 37 °C for 4 h. Formation of aggregates was verified using microscale thermophoresis (Prometheus Panta). Fibril preparations were used immediately after formation.

### Thioflavin T aggregation assays

Aggregation of HttEx1Q44 (20 µM) was monitored in 25 mM HEPES-KOH (pH 7.4), 75 mM KCl, 75 mM NaCl, and protease inhibitors following Factor Xa-mediated MBP cleavage. Thioflavin T (ThT) was added to a final concentration of 45 µM. Where indicated, FibrilPaint peptides were included at concentrations of 0.02, 0.2, or 2 µM. Fluorescence measurements were collected every 5 min for 24 h at 37 °C with orbital shaking (600 rpm) using a CLARIOstar® Plus plate reader.

### Microscale thermophoresis (MST)

Binding interactions between FibrilPaint1 and HttEx1Q44 monomers or fibrils were assessed using a Monolith MST system (NanoTemper Technologies). Measurements were performed with 50 nM fluorescently labeled FibrilPaint1 and a dilution series of HttEx1Q44 in 25 mM HEPES-KOH (pH 7.5), 75 mM KCl, and 75 mM NaCl. Samples were loaded into premium capillaries and measured at 37 °C using medium LED power and 50% MST laser power. Thermophoresis was induced 1 s after measurement start and recorded for 20 s. Signal values at 5 s were used for analysis.

### Electron microscopy

Fibril samples were diluted to final concentrations between 2 µM and 20 µM and kept on ice prior to grid preparation. Carbon-coated copper grids were glow-discharged in air (0.1 bar, 10 mA, 15 s). A 2 µL aliquot of sample was applied to the grid and incubated for 60 s, followed by blotting, washing twice with Milli-Q water, and staining twice with 2% (w/v) uranyl acetate. The second stain was incubated for 60 s before blotting and air drying.

Imaging was performed using a Talos L120C transmission electron microscope (Thermo Fisher Scientific) operating at 120 kV. Images were recorded using a CETA camera at magnifications ranging from 210× to 120,000× with defocus values between −1.0 and −2.0 µm.

Fibril length and clusters were quantified from negatively stained images using Fiji (ImageJ). Only fully visible fibrils were included.

### FibrilRuler assay: size determination by Flow-Induced Dispersion Analysis (FIDA)

Fibril size measurements were conducted using Flow-Induced Dispersion Analysis (FIDA) on a FIDA1 instrument equipped with a 480 nm excitation laser. Experiments were performed in capillary dissociation mode, with capillaries pre-equilibrated in buffer prior to sample injection. To minimize surface interactions, only the indicator solution contained fibrils.

HttEx1Q44 fibrils were diluted to 2 µM in 25 mM HEPES-KOH (pH 7.5), 75 mM KCl, 75 mM NaCl, and 0.5% pluronic, supplemented with 200 nM FibrilPaint1. Data were analyzed using FIDAbio software (v2.7) with automated baseline correction at 1 min. Multiple species were identified, and the largest detected species was used for subsequent analysis.

### Dynamic light scattering (DLS) to monitor HttEx1 fibril growth

DLS measurements were performed to obtain cumulant-derived hydrodynamic radii as a comparative measure of aggregation dynamics over time, acknowledging that heterogeneous fibril populations limit absolute size determination.

HttEx1Q44 aggregation reactions were initiated by Factor Xa-mediated cleavage of the MBP tag, and samples were analyzed by DLS at defined time points. Size distributions were extracted using Prometheus Panta data analysis software from autocorrelation functions, and apparent hydrodynamic radii (R_h__app) were calculated to track changes in aggregate populations. Only measurements that passed the built-in accuracy criteria and fell within the validated detection range of the software were included in subsequent analyses.

### Ubiquitination and proteasome assays

Expression, purification, and functional characterization of the E3 ligase CHIP, as well as reconstitution of the ubiquitin transfer cascade and proteasome degradation assays, were performed as described previously [29]. Briefly, recombinant human CHIP was expressed in E. coli with a removable N-terminal His_6_–Smt tag, purified by affinity and ion-exchange chromatography, and tag-cleaved using Ulp1 protease. Purified CHIP was assessed for purity and concentration prior to use.

Ubiquitin transfer reactions were reconstituted in vitro using fluorescein-labeled ubiquitin together with recombinant E1, E2, and CHIP in the presence or absence of ATP. For targeting amyloid fibrils, reactions additionally contained the non-fluorescent FibrilPaint variant FP20 and preformed fibrils. Ubiquitin transfer over the cascade was monitored using Flow-Induced Dispersion Analysis (FIDA), as previously established [29].

Binding interactions between FibrilPaint peptides and CHIP were evaluated by microscale thermophoresis using a Monolith system, following established protocols (Paper Y). Measurements were performed at constant peptide concentration with titrated CHIP under physiological buffer conditions.

Human 26S proteasomes were affinity-purified from HEK293 cells expressing an HTBH-tagged proteasome subunit [29]. Proteasomes were eluted by TEV protease cleavage and used immediately without freezing.

For degradation assays, ubiquitinated amyloid fibrils were incubated with freshly purified 26S proteasomes in the presence or absence of the proteasome inhibitor MG132. Proteasomal processing was monitored by dynamic light scattering, and apparent hydrodynamic radii were normalized to the time point of proteasome addition to account for sample variability, as described previously [29].

## Data analysis

All data were processed using GraphPad Prism (version 10.4.1). ThT aggregation curves and FIDA-based growth data were fitted using Boltzmann sigmoidal models to extract lag times, half-times, and plateau values. Binding curves were analyzed using a three-parameter agonist–response model to determine apparent EC_50_ values.

## Notes

### Competing Interest Statement

FAD, GM, TG, JAP, AF and SGDR are named as inventors in a patent (EP23194706, Peptides for the detection of amyloid fibril Aggregates) filed by Universiteit Utrecht Holding BV describing the peptides mentioned in this manuscript. The other authors declare no competing interests.

### Summary of Updates

Minor revisions were made to the manuscript text for clarity and consistency. Figures were updated to improve presentation and methodology was expanded.

